# OmpF Downregulation Mediated by Sigma E or OmpR Activation Confers Cefalexin Resistance in *Escherichia coli* in the Absence of Acquired β-Lactamases

**DOI:** 10.1101/2021.05.16.444350

**Authors:** Maryam Alzayn, Punyawee Dulyayangkul, Naphat Satapoomin, Kate J. Heesom, Matthew B. Avison

## Abstract

Cefalexin is a widely used 1^st^ generation cephalosporin, and resistance in *Escherichia coli* is caused by Extended-Spectrum (e.g. CTX-M) and AmpC β-lactamase production and therefore frequently coincides with 3^rd^ generation cephalosporin resistance. However, we have recently identified large numbers of *E. coli* isolates from human infections, and from cattle, where cefalexin resistance is not β-lactamase mediated. Here we show, by studying laboratory selected mutants, clinical isolates, and isolates from cattle, that OmpF porin disruption or downregulation is a major cause of cefalexin resistance in *E. coli*. Importantly, we identify multiple regulatory mutations that cause OmpF downregulation. In addition to mutation of *ompR*, already known to downregulate OmpF and OmpC porin production, we find that *rseA* mutation, which strongly activates the Sigma E regulon, greatly increasing DegP production, which degrades OmpF, OmpC and OmpA porins. Furthermore, we reveal that mutations affecting lipopolysaccharide structure, exemplified by the loss of GmhB, essential for lipopolysaccharide heptosylation, also modestly activate DegP production, resulting in OmpF degradation. Remarkably, given the critical importance attached to such systems for normal *E. coli* physiology, we find evidence for DegP-mediated OmpF downregulation, *gmhB* and *rseA* loss of function mutation in *E. coli* isolates derived from human infections. Finally, we show that these regulatory mutations enhance the ability of group 1 CTX-M β-lactamase to confer reduced carbapenem susceptibility, particularly those mutations that cause OmpC in addition to OmpF downregulation.

## Introduction

Cefalexin is a 1^st^ generation cephalosporin widely used in human, companion, and farmed animal medicine. In 2016 in Bristol, United Kingdom, and surrounding regions, (a population of 1.5 million people) 27.6 cefalexin courses were dispensed per 1000 patient population (2.8% of all dispensed items). Whilst dispensing rates had dropped by 19.5% since 2013, the proportion of *Escherichia coli* from community-origin urine samples resistant to cefalexin in this region rose from 7.06% to 8.82% (1).

Cefalexin resistance in *E. coli* is caused by hyper-production of the chromosomally-encoded class 1 cephalosporinase gene *ampC*, or acquisition of plasmid AmpC (pAmpC), or Extended Spectrum β-lactamases (ESBLs). These are also mechanisms of 3^rd^ generation cephalosporin resistance (3GC-R). We recently reported that among community-origin urinary *E. coli* from Bristol and surrounding regions collected in 2017/18, 69% of cefalexin resistant isolates were 3GC-R, suggesting that cefalexin resistance in the absence of ESBL/AmpC production is common (2). A similar observation was made when analysing faecal samples from dairy cattle in the same region, where only 30% of samples containing cefalexin resistant *E. coli* yielded 3GC-R isolates (3). Hyper-production of common acquired penicillinases such as TEM-1 and OXA-1 does not confer cefalexin resistance in *E. coli* (4). Furthermore, the involvement of efflux pump over-production, e.g. AcrAB-TolC in *E. coli* has not been reported, but OmpF porin loss is known to reduce cefalexin susceptibility (5). Indeed, early work showed cefalexin more efficiently uses OmpF than OmpC porin to enter *E. coli* (6).

One aim of the work reported here was to characterise cefalexin resistance mechanisms in *E. coli* lacking acquired β-lactamases by studying resistant mutants selected *in vitro*. A second aim was to characterise mechanisms of cefalexin resistance seen in 3GC-susceptible (3GC-S) human urinary and cattle isolates from our earlier surveillance studies (2,3). A third aim was to determine if the cefalexin resistance mechanisms identified here enhance CTX-M mediated β-lactam resistance.

## Results and Discussion

### Cefalexin resistance in *E. coli* is associated with OmpF/OmpC porin downregulation due to *ompR* mutation

One spontaneous cefalexin resistant mutant was selected from each of three *E. coli* parent strains: EC17, ATCC25922 and PSA. Cefalexin MICs against these isolates and their mutant derivatives are reported in **table 1**. In each case, to identify the possible cause of cefalexin resistance, LC-MS/MS whole-cell proteomics was performed comparing each mutant with its parent. No mutant over-produced the chromosomally-encoded AmpC β-lactamase (**Tables S1-S3**), and no promoter/attenuator sequence mutations upstream of *ampC* were identified in any of the mutants, based on WGS (**Figure S1**). The only significant (p<0.05; >2-fold) protein abundance change common to all three wild-type/mutant pairs was downregulation of OmpF porin production (**Table 2, Tables S1-S3**). There was no evidence of AcrAB-TolC efflux pump over-production in the proteomics data for mutant (**Tables S1-S3**). Despite OmpF porin downregulation, comparison of *ompF*-containing WGS contigs from wild-type/mutant pairs revealed no mutations in *ompF* or within 10 kb up- or downstream. We therefore concluded that there is a *trans*-regulatory mutation affecting OmpF abundance in each mutant.

**Table 1.** MIC (μg·ml^−1^) of cefalexin against *E. coli* isolates and mutant derivatives.

| PSA | PSA<br>(M) | ATCC25922 | ATCC25922<br>(M) | EC17 | EC17<br>(M) | ATCC25922<br>*ompF* | ATCC25922<br>*rseA* | ATCC25922<br>*gmhB* | Farm-1 | Farm-2 | UTI-1 | UTI-2 |
| --- | --- | --- | --- | --- | --- | --- | --- | --- | --- | --- | --- | --- |
| 16 | 32 | 16 | 32 | 8 | 32 | 32 | 64 | 64 | 64 | 64 | 32 | 32 |

“M” designates mutants selected for growth on cefalexin.
Shaded values represent resistant based on CLSI breakpoints (23), otherwise susceptible.

**Table 2.** LC-MS/MS proteomic comparisons of porin proteins and DegP abundance in *E. coli* isolates versus cefalexin resistant mutant derivatives.

|  | **OmpF** |  |  | **OmpC** |  |  | **OmpA** |  |  | **DegP** |  |  |
| --- | --- | --- | --- | --- | --- | --- | --- | --- | --- | --- | --- | --- |
|  | **Mean** | **SEM** | **P (WT/M)** | **Mean** | **SEM** | **P (WT/M)** | **Mean** | **SEM** | **P (WT/M)** | **Mean** | **SEM** | **P (WT/M)** |
| **PSA** | 0.23 | 0.05 |  | 5.93 | 1.00 |  | 3.24 | 0.32 |  | 0.06 | 0.01 |  |
| **PSA (M)** | 0.00 | 0.00 | 0.005 | 0.17 | 0.04 | 0.002 | 3.69 | 0.74 | >0.25 | 0.03 | 0.01 | 0.03 |
| **ATCC25922** | 1.69 | 0.10 |  | 1.66 | 0.26 |  | 5.55 | 0.41 |  | 0.08 | 0.01 |  |
| **ATCC25922 (M)** | 0.29 | 0.14 | 0.0006 | 0.24 | 0.05 | 0.003 | 1.00 | 0.17 | 0.0003 | 0.54 | 0.10 | 0.005 |
| **EC17** | 1.15 | 0.15 |  | 2.49 | 0.50 |  | 3.79 | 0.29 |  | 0.17 | 0.03 |  |
| **EC17 (M)** | 0.39 | 0.16 | 0.01 | 2.25 | 0.66 | >0.25 | 4.47 | 0.78 | >0.25 | 0.74 | 0.10 | 0.003 |

Since the two-component system OmpR/EnvZ is known to control porin gene transcription in *E. coli* (7) we searched among WGS data for mutations in the genes encoding this regulator, and a mutation was found in *ompR* in the cefalexin resistant derivative of isolate PSA, predicted to cause a Gly63Ser change in OmpR. A Gly63Val substitution in OmpR has previously been shown to cause OmpF and OmpC porin downregulation in *E. coli* (8) and proteomics confirmed that OmpC was also downregulated in the PSA-derived cefalexin resistant mutant relative to PSA, but the third major porin OmpA was not (**Table 2**). Accordingly, we conclude that OmpR mutation explains cefalexin resistance due to OmpF (and possibly OmpC) downregulation in the mutant derivative of isolate PSA. However, *ompR* and *envZ* were found to be wild-type in the other two cefalexin resistant mutants, suggesting alternative regulatory mutations.

### DegP over-production due to RseA anti-Sigma E mutation is associated with OmpF porin downregulation and cefalexin resistance in *E. coli*

Nine proteins, including OmpF, were significantly differentially regulated in the same direction in the cefalexin resistant mutants derived from isolates EC17 and ATCC25922, each relative to their parent strain. Three proteins (BamD, DegP and YgiM) were upregulated and six (NmpC, DctA, ArcA, OmpF and YhiL) were downregulated (**Tables S1,S2**). We were interested to note that one upregulated protein in both mutants was DegP (**Table 2**), which is a protease known to degrade porin proteins (9,10). Interestingly, in the PSA-derived *ompR* mutant with downregulated OmpF and OmpC, described above, DegP production was 2-fold lower than in the wild-type parent, suggesting a feedback response to porin downregulation (**Table 2**). DegP production was increased 7-fold in the ATCC25922-derived mutant and OmpF was downregulated 5.9-fold, as was OmpC (6.7-fold) and OmpA (5.6-fold) (**Table 2**); which is a typical Sigma E response (11). In the EC17-derived mutant, DegP was upregulated a more modest 4.4-fold, and here, OmpF was downregulated 2.9-fold, but OmpC and OmpA were not significantly (p<0.05) downregulated, suggesting a weaker Sigma E response (**Table 2**). This led to the suggestion that OmpC downregulation, seen in the PSA-derived and ATCC25922-derived cefalexin resistant mutants alongside OmpF downregulation (**Table 2**) is not necessary for cefalexin resistance. To confirm this, we disrupted *ompF* in ATCC25922 and found this to be sufficient for cefalexin resistance (**Table 1**). Additional downregulation of OmpC is not necessary.

Analysis of WGS data identified that the ATCC25922-derived mutant expressing a phenotype typical of a strong Sigma E response had a mutation predicted to cause a Trp33Arg mutation in RseA, which is a known Sigma E anti-sigma factor (12,13). Loss of RseA is expected to release Sigma E so that it can bind, among others, to the *degP* promoter, increasing transcription, leading to porin degradation and cefalexin resistance (11). We disrupted *rseA* in ATCC25922 and confirmed that this mutation does cause cefalexin resistance (**Table 1**).

### Perturbation of Lipopolysaccharide heptosylation due to *gmhB* mutation causes cefalexin resistance in *E. coli*

The EC17-derived cefalexin resistant mutant which also appears to have a Sigma E response, though weaker than the *rseA* mutant, was shown through WGS analysis to have a deoxythymidine nucleotide insertion after nucleotide 348 of *gmhB*, predicted to cause a frameshift affecting the encoded protein beyond amino acid 117. This gene encodes the enzyme D-alpha,beta-D-heptose-1,7-bisphosphate phosphatase, which is part of a pathway responsible for producing heptose for lipopolysaccharide biosynthesis (14). Loss of enzymes involved in this system are associated with increased outer membrane permeability, but interestingly, deletion of *gmhB* does not disrupt full length LPS production or damagingly compromise the outer membrane permeability barrier (14,15). The obvious conclusion is that this perturbation in envelope structure activates the Sigma E regulon, resulting in OmpF degradation by DegP. We disrupted *gmhB* in ATCC25922 and found that this mutation causes cefalexin resistance (**Table 1**). In the ATCC25922 background, the *rseA* and *gmhB* mutants were similar, in MIC terms, to the *ompF* mutant (**Table 1**). This further supports the conclusion that despite other porin production changes caused by *rseA* mutation and *ompR* mutation, as identified above, it is OmpF downregulation that is driving the cefalexin resistance phenotype observed in these three in vitro selected mutants.

### Loss and downregulation of OmpF in cefalexin resistant *E. coli* from cattle and humans and evidence for *rseA* and *gmhB* mutations in human clinical isolates

We chose two cefalexin resistant but 3GC-S isolates at random from our previous survey of dairy farms (3), and two from our previous survey of human urinary *E. coli* (2). Cefalexin resistance was confirmed by MIC (**Table 1**). WGS revealed disruption of *ompF* in both farm isolates: in Farm-1, a Tn*5* insertion disrupted *ompF*, truncating OmpF after amino acid 316. In Farm-2 a frameshift mutation disrupted OmpF after amino acid 96.

The *ompF* gene was intact in both human urinary isolates, which were identified by WGS as being ST131. Proteomics did, however, show significant (p<0.05) downregulation of OmpF abundance relative to ribosomal proteins compared with the control human isolate EC17 (1.43 +/−0.16, Mean +/- SEM, n=3) and a very closely phylogenetically related control ST131 urinary isolate, collected in parallel (3), UTI-80710 (1.15 +/− 0.09, n=3), in both cefalexin resistant urinary isolates. In UTI-1, OmpF downregulation was ~2-fold relative to both controls (OmpF abundance: 0.70 +/−0.12, n=3), but in UTI-2, OmpF was ~10-fold downregulated relative to both controls (OmpF abundance: 0.13 +/−0.03, n=3). Notably, UTI-1 also had a nonsense mutation at codon 82 in *ompC*. As expected, therefore, OmpC was undetectable by proteomics in UTI-1, but OmpC abundance relative to ribosomal proteins in UTI-2 (2.64 +/−0.84, n=3) was not significantly lower (p>0.25) than in control isolates UTI-80710 (3.14 +/−0.31, n=3) and EC17 (2.55 +/−0.61, n=3). Most interestingly, UTI-2 produced >2-fold elevated (p<0.05) levels of DegP (abundance relative to ribosomal proteins: 0.42 +/−0.04, n=3) compared with control control isolates EC17 (0.20 +/−0.04, n=3) and UTI-80710 (0.13 +/−0.02, n=3), suggestive of a phenotype like that of the *gmhB* mutant, described above. UTI-1 did not produce DegP at levels significantly different from control (p>0.25).

According to WGS, ST131 isolate UTI-2 did not have a mutation in *gmhB, rseA, ompR*, or *ompF* relative to the ST131 control isolate UTI-80710. Therefore, the regulatory mutation leading to elevated DegP levels, reduced OmpF levels, and cefalexin resistance in UTI-2 has not been identified. However, given the complexity of Sigma E activation signals and the impact that many different changes in envelope structure can have on it (16), it is possible that clinical isolates do carry mutations that activate this regulon. Indeed, searches of the NCBI database identified carbapenem resistant human *E. coli* isolate E300, identified in Japan (17), which has an 8 nt insertion, leading to a frameshift in *rseA* at nucleotide 34 (Accession Number AP022360). Furthermore, two human clinical isolates were found to have a single nucleotide insertion leading to a frameshift in *gmhB* after nucleotide 126; one from China (Accession Number CP008697) and one from the USA (Accession Number CP072911); and three commensal *E. coli* from the USA (18) were found to have frameshift mutations at various positions in *gmhB* (Accession Numbers CP051692, CP054319, and CP054319). Accordingly, we conclude that mutations likely to cause the same phenotypes found in our laboratory-selected cefalexin-resistant mutants are also found in clinical and commensal *E. coli* from across the world.

### Influence of *ompF* porin loss and downregulation on late generation cephalosporin and carbapenem susceptibility in the presence of various CTX-M β-lactamases

Our final aim was to test the impact of *ompF* loss and downregulation, due to OmpR mutation or activation of Sigma E, on late generation cephalosporin or carbapenem MIC in *E. coli* producing CTX-M β-lactamases. To do this we introduced, using conjugation, natural plasmids carrying various *bla*_CTX.M_ variants commonly identified in human and cattle 3GC-R *E. coli* in South West England: encoding CTX-M-1, CTX-M-14 and CTX-M-15 (2,19). We measured MICs of 3GCs and 4GCs used in humans (ceftazidime, cefepime) or cattle (ceftiofur, cefquinome), and the carbapenem ertapenem against CTX-M plasmid transconjugants of *E. coli* parent strains and their *ompF, rseA* or *ompR*, mutant derivatives (**Table 3**).

**Table 3.** Influence of *ompF, rseA* and *ompR* mutations on late generation cephalosporin and carbapenem MICs against *E. coli* producing CTX-M variants.

| Strain Name | MIC µg.ml<sup>-1</sup> |  |  |  |  |
| --- | --- | --- | --- | --- | --- |
|  | Cefepime | Cefquinome | Ceftazidime | Ceftiofur | Ertapenem |
| ATCC25922(pK18) | 0.25 | 0.125 | 0.25 | 0.25 | 0.016 |
| ATCC25922(pK18) CTX-M-1 | >128 | >128 | 16 | >128 | 0.0625 |
| ATCC25922(pK18) CTX-M-15 | >128 | >128 | 32 | >128 | 0.125 |
| ATCC25922(pK18) CTX-M-14 | 64 | >128 | 2 | >128 | 0.0313 |
| ATCC25922(pK18) *ompF* | 0.25 | 0.125 | 0.5 | 0.5 | 0.016 |
| ATCC25922(pK18) *ompF* CTX-M-1 | >128 | >128 | 32 | >128 | 0.25 |
| ATCC25922(pK18) *ompF* CTX-M-15 | >128 | >128 | 64 | >128 | 0.5 |
| ATCC25922(pK18) *ompF* CTX-M-14 | >128 | >128 | 4 | >128 | 0.0313 |
| ATCC25922(pK18) *rseA* | 0.25 | 0.25 | 0.5 | 0.5 | 0.0313 |
| ATCC25922(pK18) *rseA* CTX-M-1 | >128 | >128 | 16 | >128 | 0.5 |
| ATCC25922(pK18) *rseA* CTX-M-15 | >128 | >128 | 32 | >128 | 0.5 |
| ATCC25922(pK18) *rseA* CTX-M-14 | >128 | >128 | 4 | >128 | 0.125 |
| PSA | 0.125 | 0.125 | 0.5 | 0.25 | 0.016 |
| PSA CTX-M-1 | >128 | >128 | 32 | >128 | 0.125 |
| PSA CTX-M-15 | >128 | >128 | 32 | >128 | 0.25 |
| PSA CTX-M-14 | 128 | >128 | 8 | >128 | 0.0313 |
| PSA (M) (*ompR*) | 0.125 | 0.25 | 0.5 | 0.5 | 0.0625 |
| PSA (M) (*ompR*) CTX-M-1 | >128 | >128 | 16 | >128 | 1 |
| PSA (M) (*ompR*) CTX-M-15 | >128 | >128 | 16 | >128 | 1 |
| PSA (M) (*ompR*) CTX-M-14 | >128 | >128 | 4 | >128 | 0.5 |

CTM-M variants were delivered on natural plasmids by conjugation. Plasmid pK18 (32) was added to provide a marker (kanamycin resistance) to allow selection for recipients in conjugation. Isolate PSA is fluoroquinolone resistant, so this was not necessary. Shading is “non-susceptible” (resistant or intermediate) according to CLSI breakpoints (23).

In wild-type ATCC25922, as expected, CTX-M-1 and CTX-M-15 conferred resistance to all four cephalosporins tested, CTX-M-14 did not confer ceftazidime resistance, and none of the enzymes conferred ertapenem resistance. Disruption of *ompF* or *rseA* did not change the susceptibility profile but there were some MIC changes. Disruption of *ompF* caused a two-doubling increase in ertapenem MIC against transconjugants producing CTX-M-1 and CTX-M-15, but there was no change in MIC against the CTX-M-14 transconjugant. Disruption of *rseA* caused a similar impact on ertapenem MIC against CTX-M-1 or CTX-M-15 producers, but additionally caused a two-doubling increase in MIC against the CTX-M-14 producer. It is likely that this additional effect is due to the downregulation in OmpC additionally seen in the *rseA* mutant (**Table 2**), OmpC being a key carbapenem porin (20).

Resistance profiles seen in wild-type *E. coli* isolate PSA CTX-M plasmid transconjugants were almost identical to transconjugants of isolate ATCC25922 (**Table 3**). However, carriage of plasmids encoding CTX-M-1 or CTX-M-15 conferred ertapenem non-susceptibility in the PSA *ompR* mutant, the ertapenem MIC being one doubling higher than against CTX-M-1 or CTX-M-15 transconjugants of the ATCC25922 *rseA* mutant derivative (**Table 3**). The greater impact of *ompR* mutation than *rseA* mutation on reducing OmpC levels (**Table 2**) likely explains this difference. Indeed, *ompR* mutation has previously been associated with ertapenem non-susceptibility in ESBL producing *E. coli* (8,20).

### Conclusions

Cefalexin is a widely used antibacterial in human and veterinary medicine, and so cefalexin resistance is of considerable clinical importance. Despite this, mechanisms of resistance have not been given very much attention, particularly in the post-genomic age. We were surprised to find that, in our recent surveys of human and cattle cefalexin resistant isolates, acquired cephalosporinase (pAmpC or ESBL) or chromosomal AmpC hyper-production were not the cause of cephalexin resistance in a large proportion of isolates (2,3). We show here strong evidence that OmpF loss or downregulation is a key mechanism of cefalexin resistance in *E. coli* in the absence of β-lactamase production. Whilst OmpF loss contributes to resistance to a wide range of antibacterials (5), our findings show that cefalexin resistance is unusual in being caused solely by OmpF loss. Furthermore, we show that OmpF downregulation can also confer this phenotype. This may explain why *ompF* loss-of-function mutations are found among *E. coli* from clinical samples, but our work suggests there may also be numerous different regulatory mutations found among clinical isolates, each downregulating OmpF.

Such is the wide range of regulatory systems controlling OmpF production, both at transcriptional, translational, and post-translational levels (7, 16, 21), it is not surprising that cefalexin resistance mutations arise in many different genes in the laboratory, as seen here. Importantly, these mutations may contribute to resistance to other antibacterials, when partnered with other mechanisms. Indeed, some of these mutations also affect OmpC levels, and if this is a sufficiently large effect (e.g. in the *ompR* mutant identified here) that can give rise to carbapenem non-susceptibility if the mutant acquires a common ESBL such as CTX-M-15. Similar *ompR* mutants have been seen in the clinic (8). The other regulatory mutations affecting OmpF levels found in the laboratory-selected mutants reported here work through Sigma E mediated DegP over-production. It is well known that DegP degrades porins (10, 16), but it has not previously been reported that DegP-mediated degradation of OmpF is sufficient to cause resistance to any antibacterial drug.

Our findings are also potentially important because they suggest that OmpF is more susceptible to DegP mediated proteolysis *in vivo* in *E. coli* than the other two main porins, OmpC and OmpA. In the *rseA* mutant with “maximal” Sigma E activation and DegP upregulation, OmpF, OmpC and OmpA levels all fell, but in the *gmhB* mutant overproducing DegP to a lesser extent, only OmpF levels significantly fell (**Table 2**). This was still sufficient to cause cefalexin resistance but had a smaller effect on ertapenem MIC in the presence of CTX-M-15 production (**Table 3**), likely due to the OmpF-specific effect on porin downregulation.

It is known that mutations affecting outer membrane and lipopolysaccharide structure activate Sigma E, because they affect envelope integrity (16). It has not previously been shown, however, that mutations disrupting *gmhB* can do this, and further, that this can cause cefalexin resistance. We considered that despite such mutations arising in the laboratory, this perhaps overstates their clinical relevance, because disruption of the Gmh system causes significant attenuation and increased susceptibility to envelope stresses, though significantly, loss of GmhB has the mildest effect in this regard (15). Accordingly, we were very interested to find cefalexin resistant human urinary ST131 isolates having OmpF downregulation, and in one case, DegP upregulation, suggestive of a GmhB negative phenotype, though *gmhB* was intact and the nature of the mutation responsible will be the focus of future work. However, importantly, we did find clear evidence of *rseA* and *gmhB* loss-of-function mutations among clinical and commensal *E. coli* from secondary analysis of WGS data. Therefore, we provide here strong evidence that mutations constitutively activating Sigma E, including those which do this by altering lipopolysaccharide structure, can be tolerated by *E. coli* in a clinical setting. These mutations, and possibly others yet to be identified, cause clinically-relevant cefalexin resistance in the absence of β-lactamase production through DegP-mediated OmpF proteolysis.

## Experimental

### Bacterial isolates, selection of resistant mutants and susceptibility testing

Three β-lactam susceptible *E. coli* isolates were used: the type-strain ATCC25922; the human urinary isolate EC17, provided by Dr Mandy Wootton, Public Health Wales; and a ciprofloxacin resistant isolate, PSA, from faecal samples collected on a dairy farm (3). To select cefalexin-resistant derivatives of these isolates, 100 μl of overnight culture grown in Nutrient Broth were spread onto Mueller Hinton agar containing 16 μg.ml^−1^ cefalexin and each plate incubated for 24 h. In addition, four cefalexin resistant but 3GC-S isolates were used: one each from faecal samples from two dairy farms (Farm-1, Farm-2) as collected previously (3); two human urinary isolates (UTI-1, UTI-2) also as collected previously (2). The control isolate UTI-80710 is 3GC-R due to CTX-M-15 production (2) and was selected based on its production of wild-type OmpF porin levels (see text). Microtiter MIC assays were performed and interpreted according to CLSI guidelines (22, 23).

### Proteomics

One millilitre of an overnight Cation Adjusted Mueller Hinton Broth (CA-MHB) culture was transferred to 50 ml CA-MHB and cells were grown at 37°C to 0.6-0.8 OD_600_. Cells were pelleted by centrifugation (10 min, 4,000×g, 4°C) and resuspended in 35 ml of 30 mM Tris-HCl, pH 8 and broken by sonication using a cycle of 1 s on, 0.5 s off for 3 min at amplitude of 63% using a Sonics Vibracell VC-505TM (Sonics and Materials Inc., Newton, Connecticut, USA). The sonicated samples were centrifuged at 8,000×g for 15 min at 4°C to pellet intact cells and large cell debris. Protein concentrations in all supernatants were quantified using the Biorad Protein Assay Dye Reagent Concentrate according to the manufacturer’s instructions. Proteins (1 μg/lane) were separated by SDS-PAGE using 11% acrylamide, 0.5% bis-acrylamide (Biorad) gels and a Biorad Min-Protein Tetracell chamber model 3000X1. Gels were resolved at 200 V until the dye front had moved approximately 1 cm into the separating gel. Proteins in all gels were stained with Instant Blue (Expedeon) for 5 min and de-stained in water. LC-MS/MS data was collected as previously described (24). The raw data files were processed and quantified using Proteome Discoverer software v1.4 (Thermo Scientific) and searched against bacterial genome and horizontally acquired resistance genes as described previously (25).

### Whole genome sequencing and analyses

WGS was performed by MicrobesNG (https://microbesng.uk/) on a HiSeq 2500 instrument (Illumina, San Diego, CA, USA) using 2×250 bp paired end reads. Reads were trimmed using Trimmomatic (26) and assembled into contigs using SPAdes 3.13.0 (27). Contigs were annotated using Prokka 1.2 (28). Resistance genes and sequence types, according to the Achtman scheme (29) were assigned using the ResFinder (30) and MLST 2.0 on the Center for Genomic Epidemiology (http://www.genomicepidemiology.org/) platform. Pairwise contig alignments to identify mutations versus parent isolate, which were sequenced in parallel, was with EMBOSS Stretcher (https://www.ebi.ac.uk/Tools/psa/emboss_stretcher/)

### Insertional inactivation of genes and conjugation of CTX-M-encoding plasmids

Insertional inactivation of *ompF, rseA*, or *gmhB* was performed using the pKNOCK suicide plasmid (31). DNA fragments were amplified with Phusion High-Fidelity DNA Polymerase (NEB, UK) from *E. coli* ATCC25922 genomic DNA by using primers *ompF*-KO-FW (5′-CAA<u>GGATCC</u>TGATGGCCTGAACTTC-3’) with a BamHI restriction site, underlined, and *ompF*-KO-RV (5′-CAA<u>GTCGAC</u>TTCAGACCAGTAGCC-3’) with a SalI site; *rseA*-KO-FW (5’-CGC<u>GGATCC</u>TGCAGAAAACCAGGGAAAGC-3’) with a BamHI site and *rseA*-KO-RV (5’-TGCA<u>CTGCAG</u>CCATTTGGGTAAGCTGTGCC-3’) with a PstI site; *gmhB-KO-*FW (5’-TAT<u>ACTAGT</u>CACGGCTATGTCCATGAGA-3’) with a SpeI site, and *gmhB-*KO-RV (5’-TAT<u>GTCGAC</u>TCGGTCAGCGTTTCAAAC-3’) with a SalI site. Each PCR products were ligated into pKNOCK-GM (31) at the BamHI and SalI (for *ompF*), BamHI and PsaI (for *rseA*), or SpeI and SalI (for *gmhB)* sites. Each recombinant plasmid was then transferred by conjugation into *E. coli* ATCC25922 previously transformed to kanamycin resistance by introducing pK18 (32) by electroporation. Mutants were selected for gentamicin non-susceptibility (10 μg.ml^−1^), with kanamycin (30 μg.ml^−1^) being used to counter-select against the donor. Mutations were confirmed by PCR using primers *ompF-F* (5’-ATGATGAAGCGCAATAAT-3’) and BT543 (5’-TGACGCGTCCTCGGTAC-3’); *rseA*-F (5’-AGCCGCTATCATGGATTGTC-3’) and BT87 (5’-TGACGCGTCCTCGGTAC-3’); *gmhB-F* (5’-TAAATCAATCAGGTTTATGC-3’) and BT543.

Conjugation of natural CTX-M-encoding plasmids from cattle *E. coli* isolates (19) YYZ70-1 (CTX-M-15), YYZ16-1 (CTX-M-1) and PSA37-1 (CTX-M-14) into *E. coli* derivatives was performed by mixing on agar. Donor and recipient strains were grown overnight on LB agar plates with selection. A loopful of colonies for each was resuspended separately into 1 ml of PBS and centrifuged at 12,000xg for 1 min. The pellet was then resuspended in 1 ml 100 mM CaCl_2_ and incubated on ice for 30 min. A 3:1 v/v ratio of recipient to donor cell suspension was made and 4 μl of the mixture were spotted onto non-selective LB agar, which was incubated for 4-5 h at 37°C. Spots of mixed growth were scraped into a micro centrifuge tube containing 500 μl PBS and 30 μl of this mixture were spread onto a selective plate and incubated overnight at 37°C. The donor *E. coli* used were isolates PSA or ATCC25922, and their derivatives. PSA is resistant to ciprofloxacin, so 4 μg.ml^−1^ ciprofloxacin was used as counter-selection against the donor. For ATCC25922 and derivatives, pK18 (32) was introduced by electroporation prior to conjugation to allow counter selection using kanamycin (50 μg.ml^−1^). Selection for the transconjugant was with 10 μg.ml^−1^ cefotaxime.

## Acknowledgements

Genome sequencing was provided by MicrobesNG (http://www.microbesng.uk).

## Transparency declaration

The authors declare no conflict of interests.

## References

1. Hammond A, Stuijfzand B, Avison MB, Hay AD. 2020. Antimicrobial resistance associations with national primary care antibiotic stewardship policy: Primary care-based, multilevel analytic study. PLoS One. 15:e0232903.

2. Findlay J, Gould VC, North P, Bowker KE, Williams OM, MacGowan AP, Avison MB. 2020. Characterization of cefotaxime-resistant urinary *Escherichia coli* from primary care in South-West England 2017-18. J Antimicrob Chemother. 75:65–71.

3. Schubert H, Morley K, Puddy EF, Arbon R, Findlay J, Mounsey O, Gould VC, Vass L, Evans M, Rees GM, Barrett DC, Turner KM, Cogan TA, Avison MB, Reyher KK. 2021. Reduced Antibacterial Drug Resistance and *blaCTX.M* β-Lactamase Gene Carriage in Cattle-Associated *Escherichia coli* at Low Temperatures, at Sites Dominated by Older Animals, and on Pastureland: Implications for Surveillance. Appl Environ Microbiol. 87:e01468–20.

4. Wu PJ, Shannon K, Phillips I. 1994. Effect of hyperproduction of TEM-1 beta-lactamase on in vitro susceptibility of *Escherichia coli* to beta-lactam antibiotics. Antimicrob Agents Chemother. 38:494–498.

5. Phan K, Ferenci T. 2017. The fitness costs and trade-off shapes associated with the exclusion of nine antibiotics by OmpF porin channels. ISME J. 11:1472–1482.

6. Kobayashi Y, Takahashi I, Nakae T. 1982. Diffusion of beta-lactam antibiotics through liposome membranes containing purified porins. Antimicrob Agents Chemother. 22:775–780.

7. Pratt LA, Hsing W, Gibson KE, Silhavy TJ. 1996. From acids to *osmZ:* multiple factors influence synthesis of the OmpF and OmpC porins in *Escherichia coli*. Mol Microbiol. 20:911–7.

8. Dupont H, Choinier P, Roche D, Adiba S, Sookdeb M, Branger C, Denamur E, Mammeri H. 2017. Structural Alteration of OmpR as a Source of Ertapenem Resistance in a CTX-M-15-Producing Escherichia coli O25b:H4 Sequence Type 131 Clinical Isolate. Antimicrob Agents Chemother. 61:e00014–17.

9. Spiess C, Beil A, Ehrmann M. 1999. A temperature-dependent switch from chaperone to protease in a widely conserved heat shock protein. Cell. 97:339–47.

10. Sklar JG, Wu T, Kahne D, Silhavy TJ. 2007. Defining the roles of the periplasmic chaperones SurA, Skp, and DegP in *Escherichia coli*. Genes Dev. 21:2473–84.

11. Ades SE. 2008. Regulation by destruction: design of the sigmaE envelope stress response. Curr Opin Microbiol. 11:535–40.

12. Missiakas D, Mayer MP, Lemaire M, Georgopoulos C, Raina S. 1997. Modulation of the *Escherichia coli* sigmaE (RpoE) heat-shock transcription-factor activity by the RseA, RseB and RseC proteins. Mol Microbiol. 24:355–71.

13. De Las Peñas A, Connolly L, Gross CA. 1997. The sigmaE-mediated response to extracytoplasmic stress in *Escherichia coli* is transduced by RseA and RseB, two negative regulators of sigmaE. Mol Microbiol. 24:373–85.

14. Taylor PL, Sugiman-Marangos S, Zhang K, Valvano MA, Wright GD, Junop MS. 2010. Structural and kinetic characterization of the LPS biosynthetic enzyme D-alpha, beta-D-heptose-1,7-bisphosphate phosphatase (GmhB) from Escherichia coli. Biochemistry. 49:1033–1041.

15. Kneidinger B, Marolda C, Graninger M, Zamyatina A, McArthur F, Kosma P, Valvano MA, Messner P. 2002. Biosynthesis pathway of ADP-L-glycero-beta-D-manno-heptose in *Escherichia coli*. J Bacteriol. 2002 184:363–9.

16. Klein G, Raina S. 2019. Regulated Assembly of LPS, Its Structural Alterations and Cellular Response to LPS Defects. Int J Mol Sci. 20:356.

17. Abe R, Akeda Y, Sugawara Y, Takeuchi D, Matsumoto Y, Motooka D, Yamamoto N, Kawahara R, Tomono K, Fujino Y, Hamada S. 2020. Characterization of the Plasmidome Encoding Carbapenemase and Mechanisms for Dissemination of Carbapenem-Resistant Enterobacteriaceae. 5:e00759–20.

18. Stephens C, Arismendi T, Wright M, Hartman A, Gonzalez A, Gill M, Pandori M, Hess D. 2020. F Plasmids Are the Major Carriers of Antibiotic Resistance Genes in Human-Associated Commensal *Escherichia coli*. mSphere. 5:e00709–20.

19. Findlay J, Mounsey O, Lee WWY, Newbold N, Morley K, Schubert H, Gould VC, Cogan TA, Reyher KK, Avison MB. 2020. Molecular Epidemiology of *Escherichia coli* Producing CTX-M and pAmpC β-Lactamases from Dairy Farms Identifies a Dominant Plasmid Encoding CTX-M-32 but No Evidence for Transmission to Humans in the Same Geographical Region. Appl Environ Microbiol. 87:e01842–20.

20. Tängdén T, Adler M, Cars O, Sandegren L, Löwdin E. 2013. Frequent emergence of porin-deficient subpopulations with reduced carbapenem susceptibility in ESBL-producing *Escherichia coli* during exposure to ertapenem in an in vitro pharmacokinetic model. J Antimicrob Chemother. 68:1319–26.

21. Delihas N, Forst S. 2001. MicF: an antisense RNA gene involved in response of *Escherichia coli* to global stress factors. J Mol Biol. 313:1–12.

22. Clinical and Laboratory Standards Institute. 2015. M07-A10. Methods for dilution antimicrobial susceptibility tests for bacteria that grow aerobically; approved standard, 10th ed. Clinical and Laboratory Standards Institute, Wayne, PA.

23. Clinical and Laboratory Standards Institute. 2020. M100-S30. Performance standards for antimicrobial susceptibility testing; thirtieth informational supplement. An informational supplement for global application developed through the Clinical and Laboratory Standards Institute consensus process. Clinical and Laboratory Standards Institute, Wayne, PA.

24. Jiménez-Castellanos JC, Wan Nur Ismah WAK, Takebayashi Y, Findlay J, Schneiders T, Heesom KJ, Avison MB. 2018. Envelope proteome changes driven by RamA overproduction in Klebsiella pneumoniae that enhance acquired ß-lactam resistance. J Antimicrob Chemother 73:88–94.

25. Takebayashi Y, Wan Nur Ismah WAK, Findlay J, Heesom KJ, Zhang J, Williams OM, MacGowan AP, Avison MB. 2017. Prediction of cephalosporin and carbapenem susceptibility in multi-drug resistant Gram-negative bacteria using liquid chromatography-tandem mass spectrometry. BioRxiv https://doi.org/10.1101/138594.

26. Bolger AM, Lohse M, Usadel B. 2014. Trimmomatic: a flexible trimmer for Illumina sequence data. Bioinformatics 30:2114–20.

27. Bankevich A, Nurk S, Antipov D, Gurevich AA, Dvorkin M, Kulikov AS, Lesin VM, Nikolenko SI, Pham S, Prjibelski AD, Pyshkin AV, Sirotkin AV, Vyahhi N, Tesler G, Alekseyev MA, Pevzner PA. SPAdes: a new genome assembly algorithm and its applications to single-cell sequencing. J Comput Biol. 2012;19:455–77.

28. Seemann T. Prokka: rapid prokaryotic genome annotation. Bioinformatics. 2014;30:2068–9.

29. Wirth T, Falush D, Lan R, Colles F, Mensa P, Wieler LH, Karch H, Reeves PR, Maiden MC, Ochman H, Achtman M. 2006. Sex and virulence in *Escherichia coli:*an evolutionary perspective. Mol Microbiol. 60:1136–1151.

30. Zankari E, Hasman H, Cosentino S, Vestergaard M, Rasmussen S, Lund O, Aarestrup FM, Larsen MV. Identification of acquired antimicrobial resistance genes. J Antimicrob Chemother. 2012;67:2640–4.

31. Alexeyev MF. 1999. The pKNOCK series of broad-host-range mobilizable suicide vectors for gene knockout and targeted DNA insertion into the chromosome of Gram-negative bacteria. BioTechniques 26:824–828.

32. Pridmore RD. 1987. New and versatile cloning vectors with kanamycin-resistance marker. Gene. 56:309–12.T

