## Supplementary Information for "OmpF Downregulation Mediated by Sigma E or OmpR Activation Confers Cefalexin Resistance in *Escherichia coli* in the Absence of Acquired β-Lactamases"

**[1]School of Cellular & Molecular Medicine, University of Bristol, Bristol, UK**

**[2]Biology Department, Faculty of Science, Princess Nourah Bint Abdulrahman University, Riyadh, Saudi Arabia**

**[3]University of Bristol Proteomics Facility, Bristol, UK**

### Table S1: Proteomics data for EC17 and its cefalexin resistant mutant.

| Accession | Gene | EC17 | EC17 | EC17 | EC17 (M) | EC17 (M) | EC17 (M) | T. test | Fold change |
|---|---|---|---|---|---|---|---|---|---|
| P0A6L4 | *nanA* | 0.02 | 0.02 | 0.03 | 0 | 0 | 0 | <0.01 | <0.01 |
| P09323 | *nagE* | 0.04 | 0.06 | 0.05 | 0 | 0 | 0 | <0.01 | <0.01 |
| P24215 | *uxuA* | 0.02 | 0.03 | 0.02 | 0 | 0 | 0 | <0.01 | <0.01 |
| P08244 | *pyrF* | 0.02 | 0.05 | 0.03 | 0 | 0 | 0 | <0.01 | <0.01 |
| P0A830 | *dctA* | 0.05 | 0.13 | 0.08 | 0 | 0 | 0 | 0.01 | <0.01 |
| P06971 | *fhuA* | 0.05 | 0.08 | 0.1 | 0 | 0.02 | 0.01 | 0.01 | 0.14 |
| P06129 | *btuB* | 0.24 | 0.07 | 0.11 | 0 | 0.05 | 0.02 | 0.04 | 0.17 |
| P37617 | *zntA* | 0.01 | 0.02 | 0.03 | 0 | 0.01 | 0 | 0.03 | 0.22 |
| P0A927 | *tsx* | 0.13 | 0.15 | 0.15 | 0 | 0.07 | 0.04 | <0.01 | 0.27 |
| P21420 | *nmpC* | 2.61 | 3.18 | 3.13 | 0.7 | 1.07 | 0.67 | <0.01 | 0.27 |
| P33650 | *feoB* | 0.27 | 0.12 | 0.15 | 0 | 0.08 | 0.07 | 0.03 | 0.27 |
| P39396 | *yjiY* | 0.58 | 0.87 | 0.76 | 0.08 | 0.34 | 0.23 | <0.01 | 0.29 |
| P0A9Q1 | *arcA* | 0.94 | 0.5 | 0.59 | 0 | 0.35 | 0.31 | 0.03 | 0.33 |
| P02931 | *ompF* | 1.34 | 1.26 | 0.86 | 0.69 | 0.31 | 0.17 | 0.01 | 0.34 |
| P31554 | *lptD* | 0.05 | 0.09 | 0.06 | 0 | 0.04 | 0.02 | 0.03 | 0.34 |
| P37626 | *yhiI* | 0.03 | 0.04 | 0.04 | 0 | 0.02 | 0.02 | 0.03 | 0.37 |
| P0AFI7 | *pdxH* | 0.03 | 0.05 | 0.06 | 0 | 0.03 | 0.02 | 0.04 | 0.38 |
| P60240 | *rapA* | 0.03 | 0.05 | 0.05 | 0 | 0.02 | 0.03 | 0.04 | 0.38 |
| P0ACR9 | *mprA* | 0.08 | 0.09 | 0.08 | 0 | 0.04 | 0.05 | 0.02 | 0.4 |
| P33937 | *napA* | 0.05 | 0.06 | 0.1 | 0.03 | 0.03 | 0.04 | 0.04 | 0.42 |
| P37440 | *ucpA* | 0.1 | 0.17 | 0.14 | 0.04 | 0.08 | 0.05 | 0.01 | 0.43 |
| P0AEE5 | *mglB* | 0.1 | 0.14 | 0.12 | 0.03 | 0.09 | 0.05 | 0.01 | 0.46 |
| P0AC02 | *bamD* | 0.07 | 0.07 | 0.05 | 0.16 | 0.12 | 0.09 | 0.01 | 2.07 |
| P62620 | *ispG* | 0.07 | 0.08 | 0.07 | 0.12 | 0.22 | 0.13 | 0.03 | 2.13 |
| P0C0L7 | *proP* | 0.05 | 0.06 | 0 | 0.07 | 0.1 | 0.08 | 0.04 | 2.25 |
| P0A991 | *fbaB* | 0.07 | 0.15 | 0.13 | 0.17 | 0.56 | 0.41 | 0.04 | 3.33 |
| P0A9W9 | *yrdA* | 0.02 | 0 | 0 | 0.02 | 0.02 | 0.02 | 0.03 | 3.78 |
| P0AEW6 | *gsk* | 0 | 0 | 0.03 | 0.03 | 0.05 | 0.03 | 0.04 | 4.04 |
| P0C0V0 | *degP* | 0.12 | 0.18 | 0.21 | 0.68 | 0.94 | 0.59 | <0.01 | 4.35 |
| P76372 | *wzzB* | 0.02 | 0 | 0 | 0.01 | 0.04 | 0.03 | 0.04 | 5.39 |
| P0AFH8 | *osmY* | 0.04 | 0.05 | 0 | 0.09 | 0.28 | 0.24 | 0.02 | 7.05 |
| P21169 | *speC* | 0 | 0.01 | 0 | 0.03 | 0.05 | 0.04 | <0.01 | 22.74 |
| P0ADT8 | *ygiM* | 0 | 0 | 0 | 0.05 | 0.07 | 0.07 | <0.01 | >20 |
| P76402 | *yegP* | 0 | 0 | 0 | 0.05 | 0.05 | 0.07 | <0.01 | >20 |
| P76187 | *ydhF* | 0 | 0 | 0 | 0.02 | 0.06 | 0.04 | 0.01 | >20 |

**Table S2: Proteomics data for ATCC25922 and its cefalexin resistant mutant.**

| Accession | Gene | ATCC 25922 | ATCC 25922 | ATCC 25922 | ATCC 25922 (M) | ATCC 25922 (M) | ATCC 25922 (M) | T. test | Fold change |
|---|---|---|---|---|---|---|---|---|---|
| P0CG19 | *rph* | 0.06 | 0.07 | 0.06 | 0 | 0 | 0 | <0.01 | <0.01 |
| P13024 | *fdhE* | 0.05 | 0.04 | 0.05 | 0 | 0 | 0 | <0.01 | <0.01 |
| P29012 | *dadX* | 0.05 | 0.04 | 0.04 | 0 | 0 | 0 | <0.01 | <0.01 |
| P0A8J4 | *ybeD* | 0.04 | 0.04 | 0.04 | 0 | 0 | 0 | <0.01 | <0.01 |
| P33025 | *psuG* | 0.04 | 0.04 | 0.04 | 0 | 0 | 0 | <0.01 | <0.01 |
| P0AEB7 | *yoaB* | 0.04 | 0.04 | 0.04 | 0 | 0 | 0 | <0.01 | <0.01 |
| P23894 | *htpX* | 0.05 | 0.06 | 0.05 | 0 | 0 | 0 | <0.01 | <0.01 |
| P02943 | *lamB* | 0.07 | 0.09 | 0.08 | 0 | 0 | 0 | <0.01 | <0.01 |
| P0AG24 | *spoT* | 0.01 | 0.01 | 0.01 | 0 | 0 | 0 | <0.01 | <0.01 |
| P76372 | *wzzB* | 0.06 | 0.04 | 0.05 | 0 | 0 | 0 | <0.01 | <0.01 |
| P0ACQ4 | *oxyR* | 0.03 | 0.02 | 0.02 | 0 | 0 | 0 | <0.01 | <0.01 |
| P30871 | *ygiF* | 0.05 | 0.04 | 0.03 | 0 | 0 | 0 | <0.01 | <0.01 |
| P05042 | *fumC* | 0.03 | 0.05 | 0.05 | 0 | 0 | 0 | <0.01 | <0.01 |
| P0ADU2 | *ygiN* | 0.1 | 0.1 | 0.15 | 0 | 0 | 0 | <0.01 | <0.01 |
| P0AEN8 | *fucU* | 0.06 | 0.08 | 0.1 | 0 | 0 | 0 | <0.01 | <0.01 |
| P37665 | *yiaD* | 0.03 | 0.04 | 0.02 | 0 | 0 | 0 | <0.01 | <0.01 |
| P25553 | *aldA* | 0.11 | 0.07 | 0.07 | 0 | 0 | 0 | <0.01 | <0.01 |
| P0A951 | *speG* | 0.07 | 0.07 | 0.04 | 0 | 0 | 0 | <0.01 | <0.01 |
| P0AAG8 | *mglA* | 0.02 | 0.01 | 0.01 | 0 | 0 | 0 | <0.01 | <0.01 |
| P0ADB1 | *osmE* | 0.07 | 0.06 | 0.04 | 0 | 0 | 0 | <0.01 | <0.01 |
| P0AA53 | *qmcA* | 0.02 | 0.02 | 0.01 | 0 | 0 | 0 | <0.01 | <0.01 |
| P0A6U8 | *glgA* | 0.07 | 0.03 | 0.06 | 0 | 0 | 0 | <0.01 | <0.01 |
| P64429 | *ypfJ* | 0.01 | 0.01 | 0.01 | 0 | 0 | 0 | <0.01 | <0.01 |
| P00934 | *thrC* | 0.05 | 0.03 | 0.02 | 0 | 0 | 0 | 0.01 | <0.01 |
| P0ABQ2 | *garR* | 0.05 | 0.12 | 0.1 | 0 | 0 | 0 | 0.01 | <0.01 |
| P00561 | *thrA* | 0.04 | 0.02 | 0.05 | 0 | 0 | 0 | 0.01 | <0.01 |
| P08506 | *dacC* | 0.05 | 0.02 | 0.04 | 0 | 0 | 0 | 0.01 | <0.01 |
| P0AB91 | *aroG* | 0.05 | 0.02 | 0.03 | 0 | 0 | 0 | 0.01 | <0.01 |
| P0A6N4 | *efp* | 0.15 | 0.28 | 0.12 | 0 | 0 | 0 | 0.01 | <0.01 |
| P60757 | *hisG* | 0.07 | 0.04 | 0.03 | 0 | 0 | 0 | 0.01 | <0.01 |
| P0ADX1 | *yhfA* | 0.02 | 0.01 | 0.01 | 0 | 0 | 0 | 0.01 | <0.01 |
| P39180 | *flu* | 0.07 | 0.05 | 0.14 | 0 | 0 | 0 | 0.01 | <0.01 |
| P08192 | *folC* | 0.04 | 0.04 | 0.09 | 0 | 0 | 0 | 0.02 | <0.01 |
| P37902 | *gltI* | 0.16 | 0.13 | 0.04 | 0 | 0 | 0 | 0.02 | <0.01 |
| P33232 | *lldD* | 0.34 | 0.19 | 0.11 | 0 | 0 | 0 | 0.02 | <0.01 |
| Q46845 | *yghU* | 0.03 | 0.01 | 0.01 | 0 | 0 | 0 | 0.02 | <0.01 |
| P25516 | *acnA* | 0.05 | 0.02 | 0.02 | 0 | 0 | 0 | 0.03 | <0.01 |
| P77748 | *ydiJ* | 0.03 | 0.01 | 0.01 | 0 | 0 | 0 | 0.04 | <0.01 |
| P28903 | *nrdD* | 0.27 | 0.14 | 0.23 | 0 | 0 | 0.02 | <0.01 | 0.04 |
| P21420 | *nmpC* | 3.21 | 2.66 | 3.13 | 0.28 | 0.17 | 0.19 | <0.01 | 0.07 |
| P0A830 | *dctA* | 0.16 | 0.12 | 0.16 | 0 | 0 | 0.04 | <0.01 | 0.08 |
| P06987 | *hisB* | 0.06 | 0.05 | 0.04 | 0 | 0.01 | 0 | <0.01 | 0.08 |
| P23827 | *eco* | 0.21 | 0.12 | 0.1 | 0 | 0 | 0.04 | 0.01 | 0.09 |

| | | | | | | | | | |
|---|---|---|---|---|---|---|---|---|---|
| P0A9Q1 | *arcA* | 0.47 | 0.83 | 0.63 | 0.06 | 0.06 | 0.08 | <0.01 | 0.1 |
| P07762 | *glgB* | 0.05 | 0.05 | 0.07 | 0 | 0 | 0.02 | <0.01 | 0.11 |
| P23836 | *phoP* | 0.04 | 0.06 | 0.04 | 0 | 0 | 0.02 | 0.01 | 0.13 |
| P0ABU5 | *elbB* | 0.14 | 0.13 | 0.1 | 0 | 0.05 | 0 | <0.01 | 0.13 |
| P0ABH0 | *ftsA* | 0.04 | 0.04 | 0.05 | 0 | 0 | 0.02 | <0.01 | 0.13 |
| P0A7B1 | *ppk* | 0.03 | 0.01 | 0.03 | 0 | 0 | 0.01 | 0.02 | 0.14 |
| P04425 | *gshB* | 0.05 | 0.04 | 0.04 | 0 | 0.02 | 0 | <0.01 | 0.14 |
| P39160 | *uxuB* | 0.02 | 0.06 | 0.03 | 0 | 0 | 0.02 | 0.03 | 0.14 |
| P06996 | *ompC* | 1.64 | 1.21 | 2.12 | 0.18 | 0.21 | 0.34 | <0.01 | 0.15 |
| P0AC19 | *folX* | 0.05 | 0.08 | 0.09 | 0 | 0 | 0.03 | 0.01 | 0.15 |
| P02931 | *ompF* | 1.89 | 1.59 | 1.6 | 0.56 | 0.15 | 0.15 | <0.01 | 0.17 |
| P0A910 | *ompA* | 4.82 | 5.6 | 6.24 | 1.32 | 0.75 | 0.92 | <0.01 | 0.18 |
| P0A7J0 | *ribB* | 0.09 | 0.16 | 0.06 | 0 | 0 | 0.06 | 0.03 | 0.18 |
| P0AE12 | *amn* | 0.03 | 0.02 | 0.04 | 0 | 0.01 | 0 | 0.02 | 0.18 |
| P75823 | *ltaE* | 0.11 | 0.06 | 0.1 | 0 | 0.02 | 0.03 | 0.01 | 0.19 |
| P0AB67 | *pntB* | 0.08 | 0.05 | 0.09 | 0 | 0.02 | 0.03 | 0.01 | 0.2 |
| P0A9Z1 | *glnB* | 0.05 | 0.06 | 0.08 | 0 | 0 | 0.04 | 0.01 | 0.2 |
| P0A917 | *ompX* | 0.81 | 1.15 | 0.94 | 0.39 | 0.11 | 0.11 | <0.01 | 0.21 |
| P62517 | *mdoH* | 0.04 | 0.03 | 0.05 | 0 | 0 | 0.03 | 0.02 | 0.21 |
| P0A6W9 | *gshA* | 0.04 | 0.02 | 0.04 | 0 | 0 | 0.02 | 0.03 | 0.22 |
| P0AFL3 | *ppiA* | 0.05 | 0.06 | 0.03 | 0 | 0 | 0.03 | 0.02 | 0.22 |
| P0ABH7 | *gltA* | 2.28 | 1.37 | 0.97 | 0.23 | 0.43 | 0.38 | 0.02 | 0.22 |
| P0AC33 | *fumA* | 0.4 | 0.47 | 0.78 | 0.05 | 0.18 | 0.15 | 0.01 | 0.23 |
| P0A6J5 | *dadA* | 0.21 | 0.14 | 0.14 | 0.02 | 0.05 | 0.04 | <0.01 | 0.23 |
| P28304 | *qorA* | 0.06 | 0.04 | 0.05 | 0 | 0 | 0.03 | 0.02 | 0.23 |
| P77376 | *ydgJ* | 0.04 | 0.02 | 0.03 | 0 | 0 | 0.02 | 0.03 | 0.24 |
| P0A6V1 | *glgC* | 0.17 | 0.06 | 0.12 | 0.02 | 0.02 | 0.04 | 0.02 | 0.24 |
| P39831 | *ydfG* | 0.09 | 0.09 | 0.09 | 0 | 0 | 0.07 | 0.02 | 0.25 |
| P32131 | *hemN* | 0.02 | 0.02 | 0.01 | 0 | 0 | 0.01 | 0.03 | 0.25 |
| P09546 | *putA* | 0.27 | 0.1 | 0.15 | 0.02 | 0.06 | 0.05 | 0.03 | 0.25 |
| P0A9M5 | *gpt* | 0.07 | 0.07 | 0.05 | 0 | 0 | 0.05 | 0.03 | 0.26 |
| P0A915 | *ompW* | 1.22 | 0.59 | 0.48 | 0 | 0.33 | 0.27 | 0.04 | 0.26 |
| P0A8E1 | *ycfP* | 0.06 | 0.08 | 0.1 | 0.03 | 0.03 | 0 | 0.01 | 0.27 |
| P0A905 | *slyB* | 0.45 | 0.5 | 0.6 | 0.15 | 0.11 | 0.15 | <0.01 | 0.27 |
| P0ABI4 | *corA* | 0.06 | 0.04 | 0.06 | 0 | 0 | 0.04 | 0.03 | 0.27 |
| P0AB38 | *lpoB* | 0.04 | 0.05 | 0.03 | 0 | 0 | 0.03 | 0.04 | 0.28 |
| P27306 | *sthA* | 0.16 | 0.08 | 0.14 | 0 | 0.05 | 0.06 | 0.02 | 0.28 |
| P52697 | *pgl* | 0.1 | 0.07 | 0.07 | 0 | 0 | 0.07 | 0.04 | 0.28 |
| P0A908 | *mipA* | 0.09 | 0.17 | 0.08 | 0.05 | 0 | 0.05 | 0.04 | 0.3 |
| P0A7S3 | *rpsL* | 0.44 | 0.37 | 0.17 | 0.11 | 0.18 | 0 | 0.04 | 0.3 |
| P68767 | *pepA* | 0.17 | 0.09 | 0.15 | 0 | 0.06 | 0.07 | 0.02 | 0.31 |
| P61889 | *mdh* | 2.49 | 2.39 | 1.96 | 0.57 | 0.73 | 0.81 | <0.01 | 0.31 |
| P60560 | *guaC* | 0.14 | 0.09 | 0.15 | 0 | 0.05 | 0.06 | 0.02 | 0.31 |
| P29680 | *hemE* | 0.08 | 0.05 | 0.08 | 0 | 0.03 | 0.04 | 0.01 | 0.32 |
| P30177 | *ybiB* | 0.07 | 0.05 | 0.06 | 0.03 | 0 | 0.03 | 0.01 | 0.32 |
| P37626 | *yhiI* | 0.06 | 0.03 | 0.06 | 0.02 | 0 | 0.03 | 0.04 | 0.34 |
| P0AC41 | *sdhA* | 1.22 | 0.55 | 0.93 | 0.19 | 0.37 | 0.37 | 0.02 | 0.35 |

| | | | | | | | | | |
|---|---|---|---|---|---|---|---|---|---|
| P37903 | *uspF* | 0.28 | 0.23 | 0.17 | 0.07 | 0.09 | 0.08 | <0.01 | 0.36 |
| P0ABK5 | *cysK* | 0.48 | 0.24 | 0.27 | 0.19 | 0.1 | 0.07 | 0.03 | 0.37 |
| P17445 | *betB* | 0.12 | 0.06 | 0.09 | 0 | 0.05 | 0.05 | 0.04 | 0.37 |
| P31142 | *sseA* | 0.14 | 0.14 | 0.1 | 0 | 0.05 | 0.1 | 0.04 | 0.38 |
| P0AED0 | *uspA* | 0.23 | 0.32 | 0.23 | 0 | 0.13 | 0.17 | 0.03 | 0.38 |
| P25524 | *codA* | 0.06 | 0.03 | 0.06 | 0 | 0.03 | 0.03 | 0.03 | 0.39 |
| P07014 | *sdhB* | 0.37 | 0.48 | 0.44 | 0.18 | 0.14 | 0.18 | <0.01 | 0.39 |
| P0AGE9 | *sucD* | 1.16 | 1.3 | 1.53 | 0.49 | 0.45 | 0.62 | <0.01 | 0.39 |
| P0AG16 | *purF* | 0.25 | 0.11 | 0.22 | 0.03 | 0.11 | 0.09 | 0.04 | 0.39 |
| P15640 | *purD* | 0.17 | 0.15 | 0.23 | 0.04 | 0.06 | 0.12 | 0.01 | 0.4 |
| P24186 | *folD* | 0.11 | 0.15 | 0.17 | 0.07 | 0.07 | 0.04 | 0.01 | 0.4 |
| P0ACJ0 | *lrp* | 0.15 | 0.16 | 0.09 | 0.05 | 0.06 | 0.05 | 0.01 | 0.4 |
| P33221 | *purT* | 0.24 | 0.26 | 0.28 | 0.07 | 0.13 | 0.12 | <0.01 | 0.4 |
| P0AG90 | *secD* | 0.46 | 0.4 | 0.75 | 0.1 | 0.3 | 0.28 | 0.03 | 0.41 |
| P0AG93 | *secF* | 0.04 | 0.06 | 0.07 | 0 | 0.03 | 0.04 | 0.04 | 0.42 |
| P0AFG6 | *sucB* | 1.17 | 0.75 | 1.26 | 0.28 | 0.44 | 0.6 | 0.01 | 0.42 |
| P0ABJ1 | *cyoA* | 0.48 | 0.4 | 0.48 | 0.09 | 0.18 | 0.29 | 0.01 | 0.42 |
| P0A988 | *dnaN* | 0.09 | 0.08 | 0.07 | 0 | 0.06 | 0.04 | 0.03 | 0.43 |
| P0A6I0 | *cmk* | 0.12 | 0.11 | 0.16 | 0.05 | 0.05 | 0.07 | 0.01 | 0.44 |
| P0A912 | *pal* | 0.53 | 0.63 | 0.38 | 0.32 | 0.17 | 0.22 | 0.01 | 0.46 |
| P13029 | *katG* | 0.42 | 0.27 | 0.44 | 0.11 | 0.17 | 0.25 | 0.02 | 0.46 |
| P0ABU0 | *menB* | 0.09 | 0.06 | 0.06 | 0.03 | 0.04 | 0.04 | 0.01 | 0.47 |
| P15254 | *purL* | 0.19 | 0.11 | 0.24 | 0.05 | 0.09 | 0.11 | 0.04 | 0.47 |
| P0AEE1 | *dcrB* | 0.17 | 0.17 | 0.13 | 0.07 | 0.06 | 0.09 | <0.01 | 0.48 |
| P0ADW3 | *yhcB* | 0.17 | 0.18 | 0.18 | 0.08 | 0.07 | 0.1 | <0.01 | 0.48 |
| P0A6T5 | *folE* | 0.11 | 0.15 | 0.13 | 0.07 | 0.06 | 0.06 | <0.01 | 0.48 |
| P0A853 | *tnaA* | 0.52 | 0.49 | 0.81 | 0.14 | 0.39 | 0.34 | 0.03 | 0.48 |
| P0ABJ9 | *cydA* | 0.54 | 0.36 | 0.65 | 0.1 | 0.33 | 0.33 | 0.04 | 0.49 |
| P39177 | *uspG* | 0.39 | 0.21 | 0.32 | 0.18 | 0.14 | 0.12 | 0.03 | 0.49 |
| P0AC02 | *bamD* | 0.05 | 0.08 | 0.04 | 0.09 | 0.13 | 0.12 | 0.01 | 2.09 |
| P0A6W5 | *greA* | 0.07 | 0.07 | 0 | 0.11 | 0.13 | 0.13 | 0.02 | 2.68 |
| P42630 | *tdcG* | 0.1 | 0.02 | 0.07 | 0.12 | 0.17 | 0.23 | 0.02 | 2.74 |
| P0A7T3 | *rpsP* | 0.44 | 0.21 | 0.32 | 0.98 | 0.84 | 0.91 | <0.01 | 2.83 |
| P0A6X7 | *ihfA* | 0.06 | 0.02 | 0.02 | 0.13 | 0.08 | 0.1 | 0.01 | 2.84 |
| P07017 | *tar* | 0.1 | 0.04 | 0.05 | 0.12 | 0.19 | 0.26 | 0.02 | 2.93 |
| P07363 | *cheA* | 0.03 | 0.02 | 0.03 | 0.11 | 0.06 | 0.12 | 0.01 | 3.63 |
| P0AGE6 | *chrR* | 0.05 | 0 | 0 | 0.05 | 0.06 | 0.08 | 0.04 | 3.65 |
| P0AGF6 | *tdcB* | 0.06 | 0.05 | 0.22 | 0.26 | 0.36 | 0.62 | 0.03 | 3.75 |
| P45523 | *fkpA* | 0.12 | 0.06 | 0.07 | 0.24 | 0.32 | 0.38 | <0.01 | 3.9 |
| P0ADZ0 | *rplW* | 0.47 | 0 | 0 | 0.68 | 0.63 | 0.54 | 0.02 | 3.93 |
| P04949 | *fliC* | 1.36 | 0.35 | 0.55 | 2.51 | 3.44 | 3.98 | <0.01 | 4.4 |
| P0A6E6 | *atpC* | 0 | 0.04 | 0 | 0.09 | 0.05 | 0.06 | 0.02 | 4.71 |
| P0C0V0 | *degP* | 0.1 | 0.06 | 0.07 | 0.4 | 0.49 | 0.74 | 0.01 | 7.02 |
| P37648 | *yhjJ* | 0.01 | 0.01 | 0 | 0.03 | 0.06 | 0.07 | 0.01 | 7.04 |
| P0A800 | *rpoZ* | 0.03 | 0 | 0 | 0.06 | 0.06 | 0.09 | <0.01 | 8.16 |
| P0A9H9 | *cheZ* | 0.01 | 0 | 0 | 0.05 | 0.04 | 0.1 | 0.02 | 15.55 |
| P0ADT8 | *ygiM* | 0 | 0 | 0 | 0.08 | 0.1 | 0.1 | <0.01 | >20 |

| | | | | | | | | | |
|---|---|---|---|---|---|---|---|---|---|
| P0A7N9 | *rpmG* | 0 | 0 | 0 | 0.38 | 0.23 | 0.28 | <0.01 | >20 |

**Table S3: Proteomics data for PSA and its cefalexin resistant mutant.**

| Accession | Gene | PSA | PSA | PSA | PSA (M) | PSA (M) | PSA (M) | T. test | Fold change |
|---|---|---|---|---|---|---|---|---|---|
| P21645 | *lpxD* | 0.06 | 0.07 | 0.07 | 0 | 0 | 0 | <0.01 | <0.01 |
| P77433 | *ykgG* | 0.05 | 0.05 | 0.04 | 0 | 0 | 0 | <0.01 | <0.01 |
| P31658 | *hchA* | 0.05 | 0.07 | 0.07 | 0 | 0 | 0 | <0.01 | <0.01 |
| P69922 | *fucI* | 0.02 | 0.03 | 0.04 | 0 | 0 | 0 | <0.01 | <0.01 |
| P29680 | *hemE* | 0.03 | 0.05 | 0.05 | 0 | 0 | 0 | <0.01 | <0.01 |
| P0AAJ8 | *hybA* | 0.02 | 0.03 | 0.04 | 0 | 0 | 0 | <0.01 | <0.01 |
| P24193 | *hypE* | 0.02 | 0.02 | 0.03 | 0 | 0 | 0 | <0.01 | <0.01 |
| P02931 | *ompF* | 0.15 | 0.21 | 0.32 | 0 | 0 | 0 | 0.01 | <0.01 |
| P0A7C6 | *pepE* | 0.05 | 0.03 | 0.07 | 0 | 0 | 0 | 0.01 | <0.01 |
| P04949 | *fliC* | 0.05 | 0.07 | 0.15 | 0 | 0 | 0 | 0.02 | <0.01 |
| P06996 | *ompC* | 3.96 | 6.61 | 7.23 | 0.1 | 0.22 | 0.2 | <0.01 | 0.03 |
| P25516 | *acnA* | 0.01 | 0.01 | 0.02 | 0 | 0 | 0 | <0.01 | 0.12 |
| P08194 | *glpT* | 0.03 | 0.03 | 0.03 | 0.01 | 0 | 0 | <0.01 | 0.14 |
| P11868 | *tdcD* | 0.09 | 0.08 | 0.2 | 0 | 0.02 | 0.04 | 0.03 | 0.17 |
| P0A9G6 | *aceA* | 0.04 | 0.03 | 0.08 | 0 | 0.03 | 0 | 0.04 | 0.19 |
| P13024 | *fdhE* | 0.03 | 0.05 | 0.04 | 0 | 0.02 | 0 | 0.02 | 0.2 |
| P0AAD6 | *sdaC* | 0.12 | 0.14 | 0.17 | 0.05 | 0.05 | 0 | <0.01 | 0.23 |
| P23721 | *serC* | 0.06 | 0.04 | 0.07 | 0 | 0.04 | 0 | 0.03 | 0.23 |
| P69786 | *ptsG* | 0.02 | 0.02 | 0.02 | 0 | 0 | 0.02 | 0.02 | 0.25 |
| P08192 | *folC* | 0.03 | 0.03 | 0.02 | 0 | 0.02 | 0 | 0.03 | 0.26 |
| P27830 | *rffG* | 0.02 | 0.03 | 0.03 | 0 | 0 | 0.02 | 0.04 | 0.27 |
| P13009 | *metH* | 0.02 | 0.04 | 0.04 | 0.01 | 0.01 | 0.01 | 0.01 | 0.28 |
| P0A9K9 | *slyD* | 0.17 | 0.14 | 0.16 | 0.13 | 0 | 0 | 0.03 | 0.28 |
| P63235 | *gadC* | 0.08 | 0.12 | 0.11 | 0 | 0.04 | 0.05 | 0.01 | 0.28 |
| P0AF08 | *mrp* | 0.05 | 0.06 | 0.06 | 0.02 | 0.04 | 0 | 0.01 | 0.31 |
| P09152 | *narG* | 0.02 | 0.02 | 0.03 | 0.01 | 0 | 0.01 | 0.02 | 0.31 |
| P30744 | *sdaB* | 0.07 | 0.08 | 0.05 | 0.03 | 0.05 | 0 | 0.04 | 0.39 |
| P0DMC7 | *rcsB* | 0.07 | 0.08 | 0.08 | 0.05 | 0.04 | 0 | 0.02 | 0.39 |
| P46130 | *ybhC* | 0.08 | 0.12 | 0.14 | 0 | 0.05 | 0.08 | 0.04 | 0.4 |
| P21513 | *rne* | 0.08 | 0.09 | 0.09 | 0 | 0.04 | 0.06 | 0.03 | 0.41 |
| P0A722 | *lpxA* | 0.07 | 0.07 | 0.07 | 0.03 | 0.06 | 0 | 0.04 | 0.42 |
| P00960 | *glyQ* | 0.1 | 0.1 | 0.13 | 0.06 | 0.08 | 0 | 0.04 | 0.42 |
| P25524 | *codA* | 0.04 | 0.06 | 0.05 | 0 | 0.03 | 0.03 | 0.04 | 0.43 |
| P0A8T7 | *rpoC* | 0.41 | 0.59 | 0.62 | 0.06 | 0.29 | 0.41 | 0.04 | 0.47 |
| P0C0V0 | *degP* | 0.05 | 0.08 | 0.06 | 0.01 | 0.04 | 0.04 | 0.04 | 0.49 |
| P0AEQ3 | *glnH* | 0.1 | 0.22 | 0.16 | 0.07 | 0.08 | 0.09 | 0.03 | 0.49 |
| P69908 | *gadA* | 0.58 | 0.78 | 0.88 | 0.18 | 0.51 | 0.42 | 0.02 | 0.5 |
| P69910 | *gadB* | 0.58 | 0.78 | 0.88 | 0.18 | 0.51 | 0.42 | 0.02 | 0.5 |
| P0A817 | *metK* | 0.21 | 0.19 | 0.19 | 0.07 | 0.11 | 0.11 | <0.01 | 0.5 |
| P0ACY1 | *ydjA* | 0.02 | 0.02 | 0 | 0.04 | 0.03 | 0.03 | 0.04 | 2.26 |
| P0AEQ6 | *glnP* | 0 | 0 | 0.03 | 0.04 | 0.03 | 0.03 | 0.04 | 3.4 |
| P45464 | *lpoA* | 0.03 | 0.03 | 0.04 | 0.11 | 0.13 | 0.11 | <0.01 | 3.65 |
| P39173 | *yeaD* | 0 | 0.04 | 0 | 0.04 | 0.07 | 0.06 | 0.04 | 3.73 |
| P0A8R0 | *rraA* | 0 | 0.05 | 0 | 0.08 | 0.05 | 0.07 | 0.03 | 3.9 |

| | | | | | | | | | |
|---|---|---|---|---|---|---|---|---|---|
| P0AC19 | *folX* | 0 | 0.02 | 0 | 0.03 | 0.03 | 0.03 | 0.01 | 4.8 |
| P0ABD3 | *bfr* | 0.04 | 0 | 0 | 0.08 | 0.1 | 0.1 | <0.01 | 6.7 |
| P08660 | *lysC* | 0 | 0 | 0 | 0.01 | 0.02 | 0.02 | <0.01 | >20 |

**Figure S1: *ampC* promoter sequences of *E. coli* isolates and mutants**

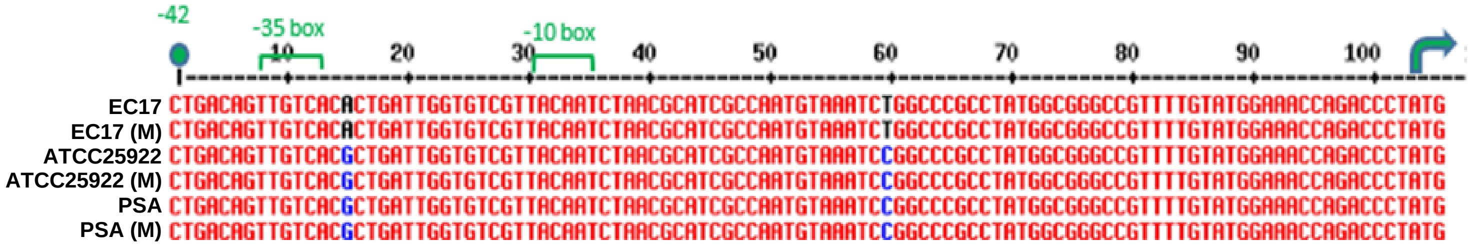
